# Network Theory Reveals Principles of Spliceosome Structure and Dynamics

**DOI:** 10.1101/2021.03.03.433650

**Authors:** Harpreet Kaur, Clarisse van der Feltz, Yichen Sun, Aaron A. Hoskins

## Abstract

Cryo-EM has revolutionized structural biology of the spliceosome and dozens of distinct spliceosome structures representing much of the splicing cycle have now been determined. However, comparison of these structures is challenging due to extreme compositional and conformational dynamics of the splicing machinery and the thousands of intermolecular interactions created or dismantled as splicing progresses. We have used network theory to quantitatively analyze the dynamic interactions of splicing factors throughout the splicing cycle by constructing structure-based networks from every protein-protein, protein-RNA, and RNA-RNA interaction found in eight different spliceosome structures. Our networks reveal that structural modules comprising the spliceosome are highly dynamic with factors oscillating between modules during each stage along with large changes in the algebraic connectivities of the networks. Overall, the spliceosome’s connectivity is focused on the active site in part due to contributions from non-globular proteins and components of the NTC. Many key components of the spliceosome including Prp8 and the U2 snRNA exhibit large shifts in both eigenvector and betweenness centralities during splicing. Other factors show transiently high betweenness centralities only at certain stages thereby suggesting mechanisms for regulating splicing by briefly bridging otherwise poorly connected network nodes. These observations provide insights into the organizing principles of spliceosome architecture and provide a framework for comparative network analysis of similarly complex and dynamic macromolecular machines.

## Introduction

The spliceosome recognizes and removes non-coding regions (introns) from precursor-mRNAs (pre-mRNAs) while ligating together the flanking exons. The spliceosome is a ribonucleoprotein (RNP), comprised of five non-coding small nuclear RNAs (snRNAs) and nearly a hundred protein factors (Wahl et al., 2009). The U1, U2, U4, U5, and U6 snRNAs in the spliceosome have distinct functions and assemble on pre-mRNAs as part of small nuclear ribonucleoprotein (snRNP) subcomplexes (the U1 and U2 snRNPs and U4/U6.U5 tri-snRNP). In addition, spliceosomes also contain a number of non-snRNP associated splicing factors including the NineTeen complex (NTC) and NTC-associated (NTA) proteins. Pre-mRNA splicing is a highly dynamic process involving assembly of splicing factors on pre-mRNAs (to form B complex spliceosomes); an activation phase in which the nascent spliceosome is remodeled to create an RNA active site (B^act^, B*1 complexes); organization and re-organization of the machinery to carry out the two sequential transesterification reactions (B*2, C, C* complexes); and finally release of the spliced mRNA from the product (P) complex and disassembly of the intron-lariat spliceosome (ILS) (Wahl et al., 2009). The entire process is driven forward by eight ATPases, each functioning at specific stages in splicing (Wahl et al., 2009).

Spliceosomes are highly dynamic in terms of both their composition and structure. More than a dozen distinct spliceosome complexes have been characterized, and it is certain that several other intermediates exist along the splicing pathway (Plaschka et al., 2019). Cryogenic electron microscopy (cryo-EM) has revolutionized studies of splicing by allowing structures of entire spliceosomes to be determined. More than 35 spliceosome structures from *Saccharomyces cerevisiae* (hereafter, yeast), *Schizosacharomyces pombe*, and *H. sapiens* have provided unparalleled insights into how splicing factors interact and function (Kastner et al., 2019; Plaschka et al., 2019; Yan et al., 2019). Nevertheless, the complexity and highly dynamic nature of the spliceosome makes comparison of different structures challenging since hundreds or thousands of intermolecular interactions may be changing from one state to the next.

Network theory provides a framework for analysis of large and complex sets of interactions such as those typified by the spliceosome. Analysis of networks provides quantitative information on the individual connectivity of each component as well as the structure of the network as a whole. At the component level, centrality parameters, such as eigenvector and betweenness centralities, can be determined for each network node. These parameters incorporate information about how that component contributes to the entire network organization. For the whole network, descriptors can include the average degree (the average number of connections for components), the algebraic connectivity (a measure of interconnectivity of all network components), and modularity (subdivisions of the network containing high connectivity between components but less to other subdivisions).

Protein-protein interaction (PPI) network analysis has been applied to many systems and is enabled by a substantial number of PPIs catalogued in various databases (MINT, STRING, DIP, PSI-MI, and InACT). These applications have broadened the scope of spectral graph theory to the understanding the biological systems, identifying the functions of novel genes, and drug discovery (Athanasios et al., 2017; Brun et al., 2003; Hasan et al., 2012; Miryala et al., 2018; Yu et al., 2013). In applications to structural biology, network analysis has been used to determine the inter-component interactions (edges) of individual residues, domains or whole molecules (nodes). These methods have been used extensively to understand the structure, folding, and function of proteins including splicing factors by analysis of different centrality metrics (Brinda and Vishveshwara, 2005; David-Eden and Mandel-Gutfreund, 2008; Fanelli et al., 2016; Menichetti et al., 2016; Negre et al., 2018; Shao et al., 2020a; Vendruscolo et al., 2002; Yan et al., 2018; Yan et al., 2014). Structural network analysis has been more limited for macromolecular machines of comparable complexity to the spliceosome, although it has been used effectively to study the peptidyl transferase center of the ribosome and intermolecular interactions observed in trypanosome mitochondrial ribosomes (David-Eden and Mandel-Gutfreund, 2008; Ramrath et al., 2018).

For the splicing machinery, network theory has been applied to both functional data from biological experiments as well as PPIs from the STRING database (Carbonell et al., 2019; Guimarães et al., 2018; Papasaikas et al., 2015; Pires et al., 2015). However, these studies are difficult to interpret with respect to molecular mechanisms since snRNAs are not part of the STRING database and without structural data the intermolecular contacts that cause changes in biological function are unclear. Recently, network theory analysis of a spliceosome structure combined with molecular dynamic simulations were used to identify putative information exchange pathways among splicing machinery components (Clf1, Cwc2, and Prp8) in the yeast C complex spliceosome (Saltalamacchia et al., 2020; Shao et al., 2020b). It is not clear, though, how the C complex structural network is assembled or changes during splicing.

Here, we calculated the structural networks of eight different spliceosome models determined from cryo-EM data representing the B➔B^act^➔B*1➔B*2➔C➔C*➔P➔ILS complexes (Fica et al., 2017; Galej et al., 2016; Plaschka et al., 2017; Wan et al., 2019; Wan et al., 2017; Wilkinson et al., 2017; Yan et al., 2016). This was enabled by creation of software that permits analysis of both RNA and protein contributions to networks using available PDB models. Comparison of these networks reveals dynamic changes in connectivity, modularity, module composition, and centrality parameters for splicing factors throughout the reaction. They also reveal how connectivity of the spliceosome becomes focused on the active site and the contributions of elongated (non-globular) and nonessential proteins make to this organization. In comparison with networks derived from PPI or functional data, our analysis has implications for understanding how perturbations propagate through different modules to influence multiple steps in splicing, how factors with high betweenness influence specific steps in splicing, and how network topological parameters correlate with lethality phenotypes.

## Results

We used network analysis to study eight models of yeast spliceosomes built from cryo-EM maps of complexes captured during activation, catalysis, and disassembly (Fica et al., 2017; Galej et al., 2016; Plaschka et al., 2017; Wan et al., 2019; Wan et al., 2017; Wilkinson et al., 2017; Yan et al., 2016). Each model was treated as its own network and analyzed individually (**Fig. 1**). First, network nodes were defined as individual single chains in the models except for proteins making up the heteroheptameric Lsm and Sm rings, which were grouped together as single nodes (*e.g*., the U2 Sm ring was considered a single node), the Prp19 homotetramer was also grouped as a single node, and the pre-mRNA substrate was divided based on continuously resolved regions in the EM density (*e.g*., split into 5’ SS and BS nodes when appropriate). The published models were further edited by assigning or removing protein chains of unknown identity and by substitution of alanine amino acids for glycines in regions where the maps precluded modeling of amino acid side chains (**SI Appendix Table S1**, **SI Appendix Discussion S1**).

**Figure 1.**
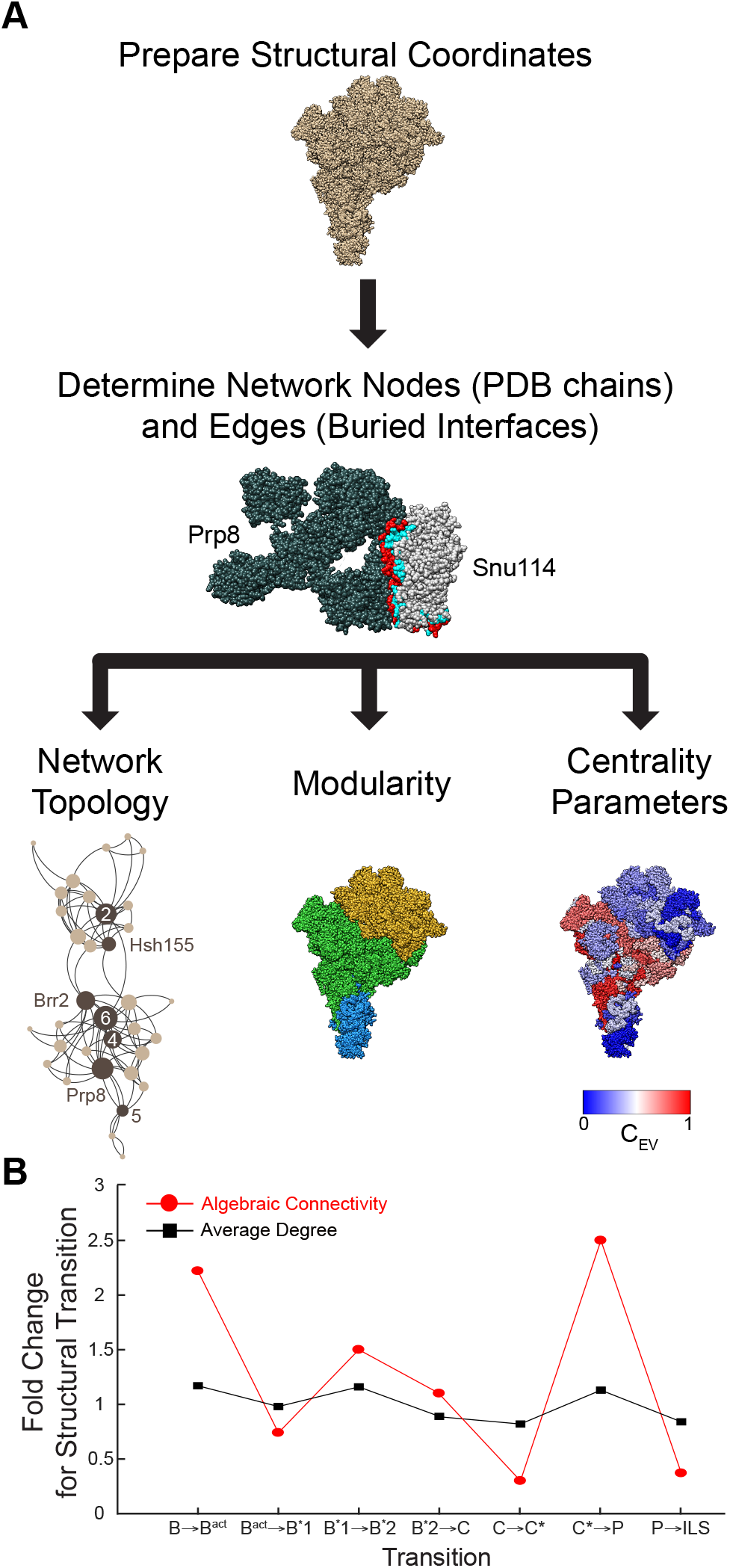
Structural network analysis workflow and network properties. **(A)**Example workflow using the yeast B complex spliceosome (shown in tan, PDB ID: 5NRL). PDB coordinate files are first edited so that network nodes and edges can be identified. Edges are calculated as the buried surface areas of molecular interfaces (*e.g*., the interface between the B complex components Prp8 (dark grey) and Snu114 (light grey) is shown in red and blue). Nodes and edges are then used to construct the network. *Bottom Left*.The network topology for B complex is shown with snRNAs (2, 4, 5, and 6) and some of the largest proteins noted (Prp8, Brr2, and Hsh155). *Bottom Middle*. Computationally defined modules in B complex are separately colored (U5 module, blue; U6 module, green; U2 module, yellow). *Bottom Right*. B complex components are colored by eigenvector centrality (C_EV_). (**B**) Fold changes in structural network parameters for spliceosomes transitioning between complexes. Structure images were generated using UCSF Chimera, and the network display was generated with Gephi.

We then defined edges in our networks as the interactions between splicing factors occuring within 1.4 Å (**Fig. 1A**). PDBePISA was used to identify these interactions from each spliceosome model (Krissinel and Henrick, 2007). The nodes and edges were then used to generate a network for each splicing complex. The network was not weighted by the amount of buried surface area (BSA) since the importance of a component’s interactions may not be directly correlated with this value and calculations of BSA in regions for which the cryo-EM maps are poor may not be accurate.

We note that two caveats of this approach are that it is restricted to models built from interpretable EM density and it assumes EM densities were interpreted correctly when models were constructed. We were unable to include information in our networks for the large amount of protein mass that is unaccounted for in all available spliceosome models (ranging from 22-38% of total amino acids, **SI Appendix Table S1**). We assume these represent protein domains which interact weakly or on the periphery of the spliceosome. We accounted for remaining uncertainties where possible by comparison of independently determined structures of spliceosomes captured in similar states to verify model accuracy (**SI Appendix Discussion S1**). While it is possible that our networks are biased towards stable interactions over those that occur transiently as well as towards interactions present in trapped spliceosomes amenable to structural elucidation, we believe they are accurate descriptions of the pre-dominant intermolecular interactions occurring within and between each spliceosome complex.

### Topology Analysis Reveals Oscillations in Algebraic Connectivity

Once the networks were computed, we first analyzed their topological information (**Fig. 1A**, **B**). The average degree (or number of connections a node makes to other nodes) ranges from 6.28-8.50 for each spliceosome network, a variation of ~1.4-fold from B to ILS complex (**Fig. 1B**, **SI Appendix Table S2**). However, the level of intermolecular connectedness of each spliceosome network (the algebraic connectivity) varies by a larger factor both for the overall transition from B to ILS complex (~2.9-fold) and for intermediary steps (**Fig. 1B**). In random networks, an increase in the number of nodes results in a decrease in algebraic connectivity. Therefore, spliceosome network topologies indicate that they are not organized randomly (*e.g*., nodes increase from 34 to 40 between the B and B^act^ networks while the connectivity doubles from 0.36 to 0.80, **SI Appendix Table S2**). This confirms that our networks capture the specific, non-random intermolecular interactions formed by biomolecular structures.

Networks computed from models of spliceosomes captured just before or after the catalytic steps of splicing (B*2, C, P) had the highest values of algebraic connectivity (**Fig. 1B**). High values of algebraic connectivity indicate a greater degree of interconnection and more difficulty in separating the network into individual components or sub-networks (Jamakovic and Van Mieghem, 2008). This shows that B*2, C, and P complex networks are the most robust and resistant to change due to removal of a given node or edge.

In contrast we observed low values of connectivity associated with the B and ILS complex networks and networks obtained from complexes transitioning to or between catalytic conformations (B*1, C*; **Fig. 1B**). The algebraic connectivity thus indicates the spliceosome is at its most interconnected during catalysis and its least interconnected while transitioning and prior to disassembly. It is interesting to note that this results in oscillations of connectivity during splicing as low connectivity states bridge those of higher connection. For example, during the C➔C*➔PàILS complex transitions, the network connectivities oscillate from 0.95➔0.35➔0.89➔0.33 while the numbers of nodes and edges remain similar (**SI Appendix Table S2**). This is consistent with a model in which formation of intermediary, low-connectivity states are required for transition between or away from structures of high connectivity.

### Spliceosome Modules are Highly Integrated and Dynamic

Spliceosomes are built from discrete complexes (the U snRNPs, NTC, and other protein splicing factors) that associate with one another and the pre-mRNA. We next studied if networks describing assembled spliceosomes maintain these complexes as discrete subnetworks that are more highly coordinated between their members than to other nodes. To do this, we calculated the modularity of each network as values ranging from 0 to a highly interconnected network to 1 for a highly modular one (6). We also calculated the number and composition of the modules for each network using MODULAR (Marquitti et al., 2014). Consistent with expectations from biochemical and structural studies, the spliceosome networks begin as highly modular in B and B^act^ complexes but then modularity decreases as the original subcomplexes integrate to form the active site (**Fig. 2A**).

**Figure 2.**
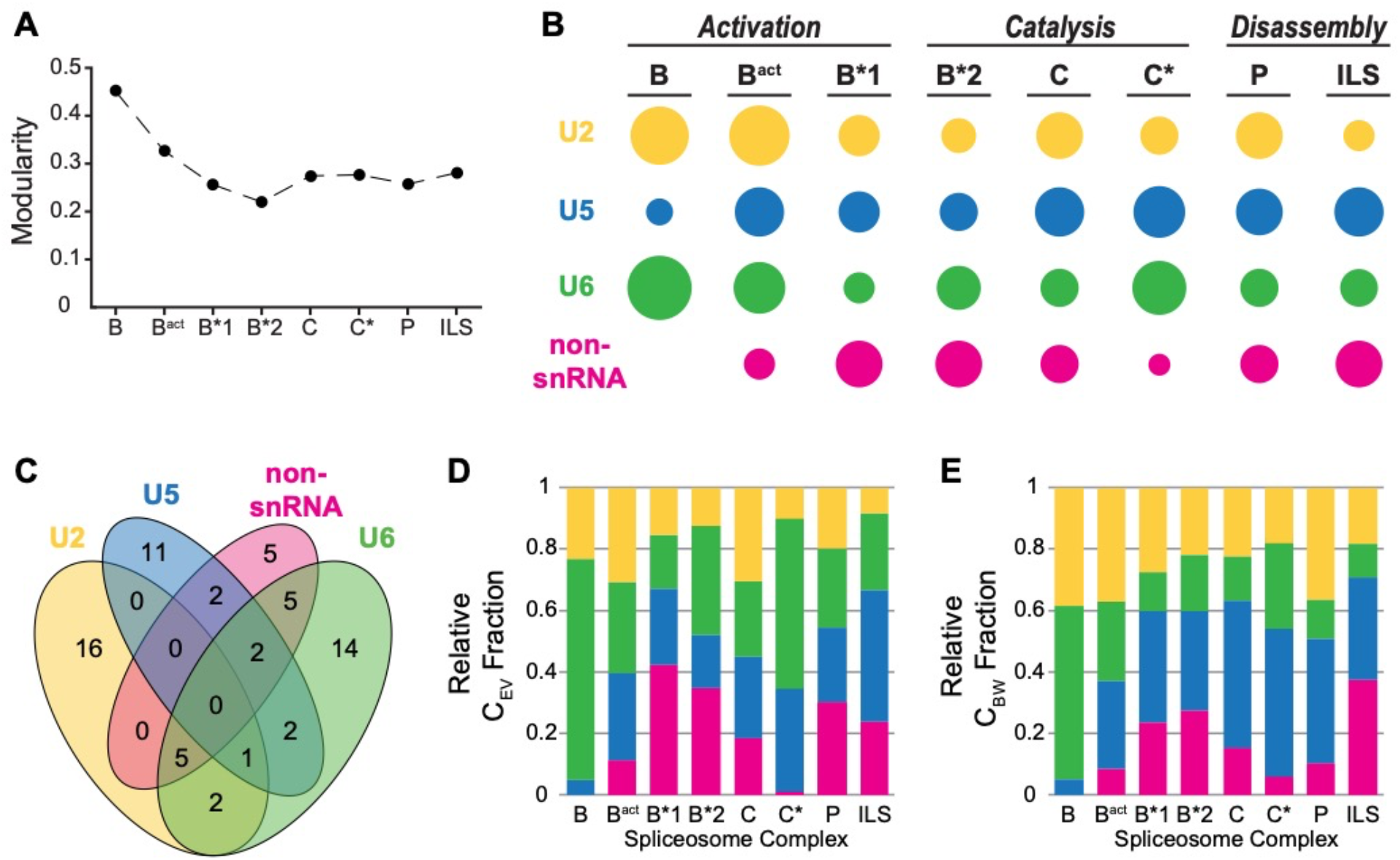
Module Integration and Component Switching during Splicing. (**A**) Calculated modularity values for spliceosome structural networks. (**B**) Schematic representation of relative module composition for each structural network. Modules are defined based on their snRNA component (U2, U5, or U6) or the absence of snRNAs (non-snRNA). The area of each circle is proportional to the number of components found in each module. Module components are listed in **SI Appendix Table S3**. (**C**) Venn diagram of the total number of protein and snRNA components in the combined BàILS structural networks associated with a given module or set of modules. (**D**) The relative contribution of each module to total C_EV_ for each structural network. (**E**) The relative contribution of each module to total C_BW_ for each structural network. For panels (B-E), colors correspond to the module nomenclature in (B).

While the overall modularity of the spliceosome networks remains nearly constant following activation, the composition of the modules is highly dynamic and can change dramatically from one structural network to the next. We categorized each module by its component snRNA (U2, U5, U6) or by the lack of snRNAs (non-snRNA). In the B complex, the modular organization of the spliceosome loosely follows the traditional description of snRNPs with three modules, each associated with a snRNA (**Fig 2B**). The U4 snRNA and snRNP proteins are associated with the U6 module in the B complex, in agreement with U4/U6 hybridization in the tri-snRNP. After the B complex, the U2, U5, and U6 snRNAs remain in distinct modules from each other and only 4 modules are needed to optimally describe the spliceosome network. As shown in in **Fig. 2B** and **SI Appendix Tables S3A-D**, the number of components in each module changes during each stage of splicing. These changes are highly variable: some components never switch modules (*e.g*., Lea1 is always present in the U2 module and Snu114 in the U5 module), others switch by themselves between modules (*e.g*., Prp8 and Brr2 switch from the U6 module to the U5 and U2 modules, respectively, during the BàB^act^ transition), and others always switch between modules as part of a group of factors (*e.g*., Cwc2 and Bud31 switch from the U6 to the non-snRNA module and back again during the B*2àCàC* transitions). The pre-mRNA substrate nodes group with snRNA-containing modules and never in the non-snRNA module. The movement of the substrate between modules reflects the catalytic steps of the splicing reaction. In B^act^ and B*1 complex networks, the 5’ SS and BS join with the U5 and U2 modules, respectively, but come together in the U6 module just prior to 5’ SS cleavage in the B*2 complex network (**SI Appendix Tables S3A-C)**. Nineteen factors in our analysis transiently associate with more than one module and eight of these associate with three different modules over the course of splicing (**Fig. 2C**, **SI Appendix Tables S3A-D**).

We were surprised to observe that the NTC/NTA complex never groups as a single module but instead spreads out over all four modules from the time of its integration in B^act^ through the ILS complex (**SI Appendix Fig. S1A**). Three of the core components of NTC/NTA (Cef1, Prp19, and Snt309) coordinately switch modules five times throughout the cycle by transiently becoming members of the U2, U6, or non-snRNA modules. Other NTC/NTA components are less dynamic and are predominantly associated with a single module (*e.g*., Cwc2, Prp17, and Bud31 are always found in the U6 module except in C complex). These results highlight the high degree of integration of the NTC/NTA into snRNA-defined modules as well as the heterogeneity of NTC/NTA component network properties. It is possible that the heterogeneity we observe in this network analysis correlates with physical or functional subcomplexes of the NTC/NTA.

One consequence of these dynamics is that while a high degree of interconnectedness and redundancy may insulate a particular network from perturbation (such as removal of an edge or node) (Guimarães et al., 2018), other structural networks along the splicing pathway may be impacted. To illustrate this concept, we deleted the node corresponding to the non-essential NTC/NTA splicing factor Ecm2 and calculated the resulting changes in connectivity in each network (**SI Appendix Fig. S1B**). The impact of Ecm2 deletion ranges from a loss of ~10% of network paths in B^act^ complex to a loss of over 40% of paths in B*1 complex. Interestingly, Ecm2 was recently proposed to play roles in both catalytic stages of splicing (van der Feltz et al., 2021). Consistent with these roles, it is in some of the spliceosome networks from these stages in which Ecm2 deletion has the greatest impact (B*1, B*2, C*, and P complexes). Since yeast cells in which Ecm2 has been deleted are viable and can splice efficiently at 30°C (van der Feltz et al., 2021; Xu et al., 1996), splicing activity is nonetheless resilient at permissive temperatures to dramatic changes in spliceosome network topology. At non-permissive temperatures (37°C), Ecm2 deletion results in a strong growth defect in yeast and a number of synthetic lethal genetic interactions 30°C (van der Feltz et al., 2021; Xu et al., 1996). How spliceosome structures and the associated networks change as a function of temperature is unknown.

### U5 and U6-snRNA Containing Modules are Major Contributors to Network Topology

We next analyzed the contribution each module makes to the overall sum of centrality parameters for each network. We computed contributions to eigenvector centrality (C_EV_, high C_EV_ indicates a major network intersection as it connects to other nodes of high connectivity; **Fig. 2D**) and betweenness centrality (C_BW_, high C_BW_ indicates a major bridge in the shortest path between nodes; **Fig. 2E**). In all networks, the snRNA-containing modules make the largest contributions to C_EV_ and C_BW_. The U5 and U6 modules are often the highest contributors to C_EV_. For U5, this echoes the large number of connections the Prp8 protein makes with splicing factors in other modules. For U6, this is consistent with its role in forming the spliceosome active site and its scaffolding by splicing factors. U6 is also the highest contributor to both C_EV_ and C_BW_ in the B complex spliceosome, reflecting the fact that interactions with and through U6 are central for the allosteric cascade of conformational changes driving spliceosome activation (Brow, 2002). It is interesting to note that while the U6 module often contributes the most highly to C_EV_, this does not always correlate with a high contribution to C_BW_ (**Fig. 2D** vs. **2E**). In other words, while many splicing factors connect to U6 within the network, many of these interactions do not result in linkages between splicing factors that are uniquely bridged by U6. This suggests a large degree of redundancy in the U6 interaction network. In contrast, the U5 module contributes very highly to C_BW_, again driven by a large number of connections that are uniquely bridged by the Prp8 protein.

### Dynamic and Step-Specific Bridges Between Splicing Factors

The high contributions by the Prp8 protein to U5 module C_EV_ and C_BW_ are consistent with its central role in spliceosome structure and its large size (Plaschka et al., 2019; Saltalamacchia et al., 2020). Prp8 contributes highly to spliceosome network centrality parameters throughout the entire reaction. However, it is not the only splicing factor to do so. The U2 snRNA and NTC/NTA component Cef1 function alongside Prp8 as the three strongest contributors to C_BW_ in the B^act^ through ILS networks (**Fig. 3A-C**).

**Figure 3.**
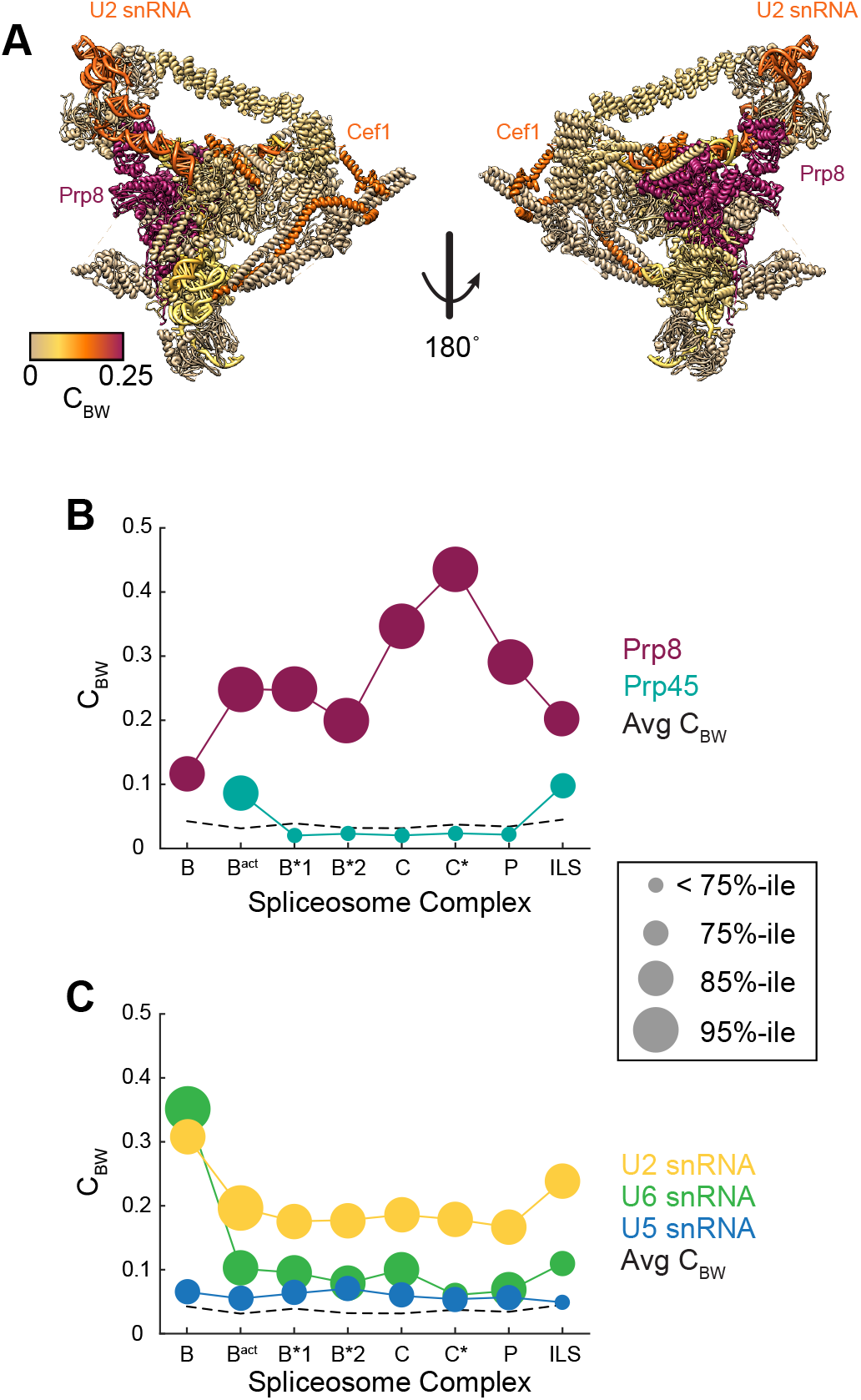
Dynamic Changes in Betweenness of Splicing Factors. (**A**) Depiction of the spliceosome B*1 complex (PDB ID: 6J6H) with components colored by C_BW_. Factors discussed in the text with high C_BW_ are identified. (**B**) Calculated C_BW_ values for Prp8 and Prp45 are plotted for each splicing factor structural network in comparison with the average C_BW_ of all splicing factors in the corresponding complex. (**C**) C_BW_ values for the snRNAs in each structural network in comparison with the average C_BW_ of all splicing factors in the corresponding complex. For (**B**) and (**C**), the circle size represents the ranking of the C_BW_ value of that component in relation to others in the complex.

Even for Prp8, structural dynamics of the spliceosome result in large changes in C_BW_ from one network to the next (**Fig. 3B**). While Prp8 is always among the components with highest C_BW_, this value changes by more than 4-fold during splicing. This results from the many interactions between nodes that are uniquely bridged by Prp8 fluctuating during splicing: conformational changes result in some of these interactions being lost or gained or no longer being unique. For example, the increase in C_BW_ for Prp8 in C and C* networks can be partially attributed to unique connections between the Prp8 and Prp16 or Prp22 ATPase nodes, respectively. The U6 snRNA is an extreme example of fluctuation in C_BW_.It is among the highest contributors to C_BW_ in the B complex network before activation results in a loss of C_BW_ due to increased path redundancy between nodes connected through U6 for the remainder of the splicing reaction (**Fig. 3C**).

Other factors can exhibit high C_BW_ only at certain stages. For example, the NTC/NTA protein Prp45 node has high C_BW_ only in B^act^ and ILS networks (**Fig. 3B**). At other stages it does not deviate from the average splicing factor C_BW_. The high values for C_BW_ in these networks are due to Prp45 serving as the major bridge for linkage of the REtention and Splicing (RES) or the NTC-Related (NTR) complexes to the center of the spliceosome (**SI Appendix Fig. S2**). Changes in C_BW_ for Prp45 are correlated with a change in the number of connections (degree) Prp45 makes to other nodes (**SI Appendix Table S4**). However, changes in degree and C_BW_ are not always correlated for splicing factors, and we identified multiple examples in which these values are also anti-correlated or independent of one another (**SI Appendix Table S4**). This shows that connectivity of the spliceosome networks does not increase or decrease in complexity simply due to the addition or subtraction of various nodes. Rather, connectivity is also changing due to rearrangement of connections between existing nodes.

### Network Interactions Focus on the Active Site

We next analyzed how the connectivity of spliceosome protein nodes corresponds with the spatial position of those same proteins relative to the active site. We approximated the spliceosome active site as being centered around the U6 snRNA nucleotide G60 and then summed C_EV_ values for all proteins that are at least partly contained within shells of increasing radius (**Fig. 4A**). This analysis shows that the proteins closest to the active site (within 10 Å) have a much higher total C_EV_ than those that located on the periphery. Interestingly, this analysis also showed that the spliceosome’s region of highest C_EV_ is the active site and not always the center of mass (**SI Appendix Table S5, Fig. S3**). Thus, the high centrality values likely reflect an abundance of functional interactions being directed towards the active site rather than being due to the amount of protein present at that location.

**Figure 4.**
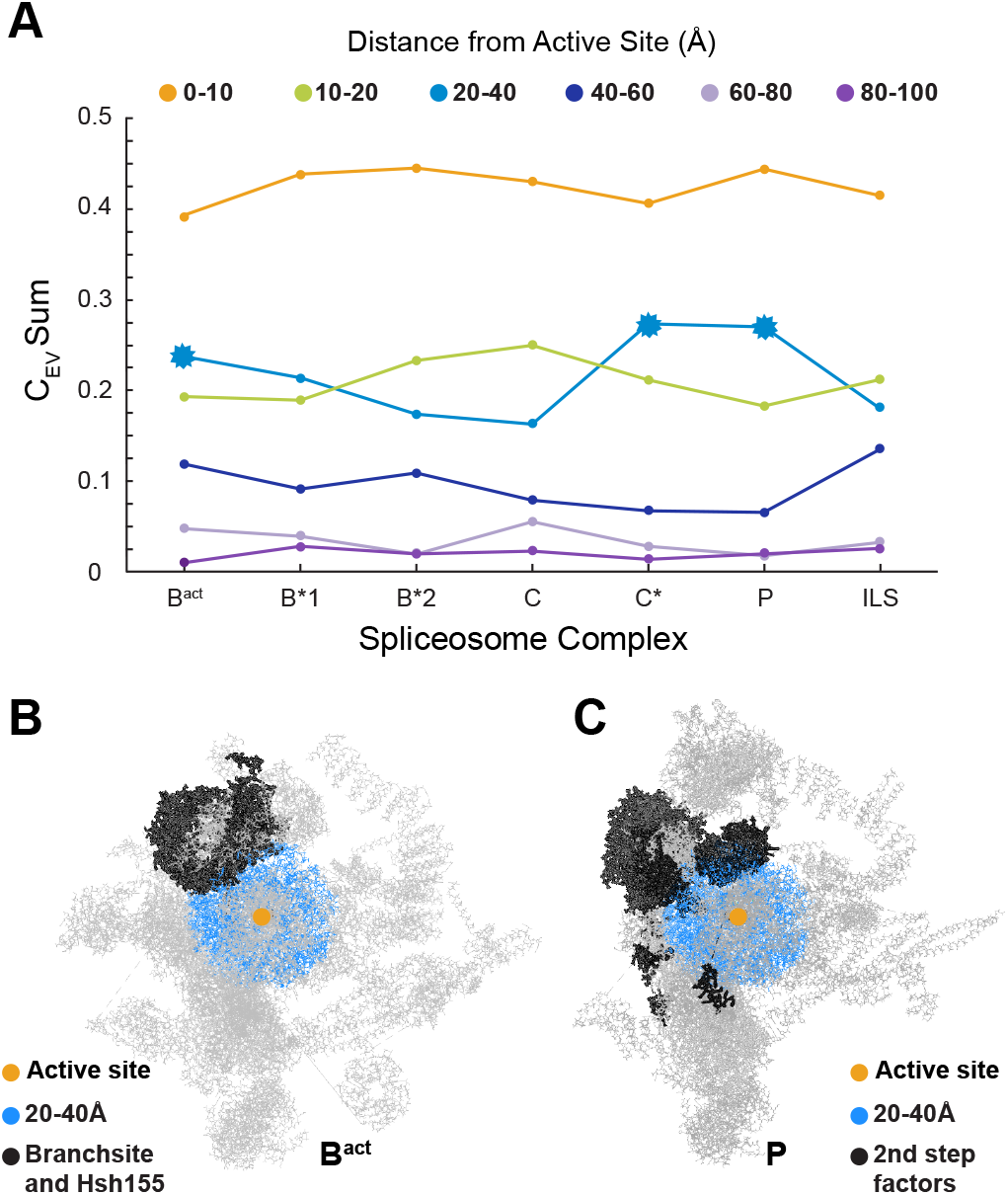
Eigenvector Centrality is Centered on the Spliceosome Active Site. (**A**) Sum of splicing factor C_EV_ values as a function of node distance from the spliceosome active site (approximated as U6 snRNA nucleotide G60). (**B**) B^act^ complex (PDB ID: 5GM6) highlighting the location of key factors in contact with the 20-40 Å shell from the active site. (**C**) P complex (PDB ID: 6EXN) highlighting the location of 2^nd^ step factors in contact with the 20-40 Å shell from the active site.

The sum of C_EV_ values decreased with increasing distance from the active site (**Fig. 4A**) indicating that the factors on the periphery of the spliceosome make fewer connections with highly interconnected components than those nearer the active site. This observation held true except for a few instances: proteins in the 20-40 Å shell had higher total C_EV_ than those in the 10-20 Å shell in B^act^, C*, and P complexes. This expansion of C_EV_ is not just due to new factors arriving but arises from the location and connection of factors in the network. Specifically, we realized the higher C_EV_ sums were due to Hsh155 (B^act^ network) and the 2^nd^ step protein factors (Prp17, Prp22, and Slu7; C* and P networks) being partially located within these shells (**Fig. 4B, C**). These proteins serve as major interaction hubs and are highly connected to splicing factors distant from the active site.

### Elongated and Nonessential Factors are Major Contributors to Connectivity

When we mapped C_EV_ values onto the proteins closest to the active site (**Fig. 5A**), we noticed that several of the factors with the highest C_EV_ shared common features: they have non-globular structures and they are conditionally non-essential (*i.e*., not required for yeast proliferation under permissive conditions) components of the yeast NTC/NTA complex. We first analyzed how non-globular proteins contribute to our calculated spliceosome networks. While many of the spliceosome’s largest proteins have well-defined three-dimensional folds (Prp8, Brr2, Hsh155), a number of proteins lack globular structural domains. We calculated surface areas for each spliceosome protein and divided these values by the number of amino acids (aa) resolved in each protein to yield the protein’s elongation factor (EF, **SI Appendix Table S6**). The distribution of EF values for splicing factors showed multiple peaks consistent with globular (*e.g*., EF^Snu114^ = ~47 Å^2^/aa), intermediate (*e.g*., EF^Cwc2^ = ~60 Å^2^/aa) and highly elongated proteins (*e.g*., EF^Isy1^ >80 Å^2^/aa) (**SI Appendix Fig. S4**). Based on this distribution, we defined elongated proteins as those with EF > 75 Å^2^/amino acid.

**Figure 5.**
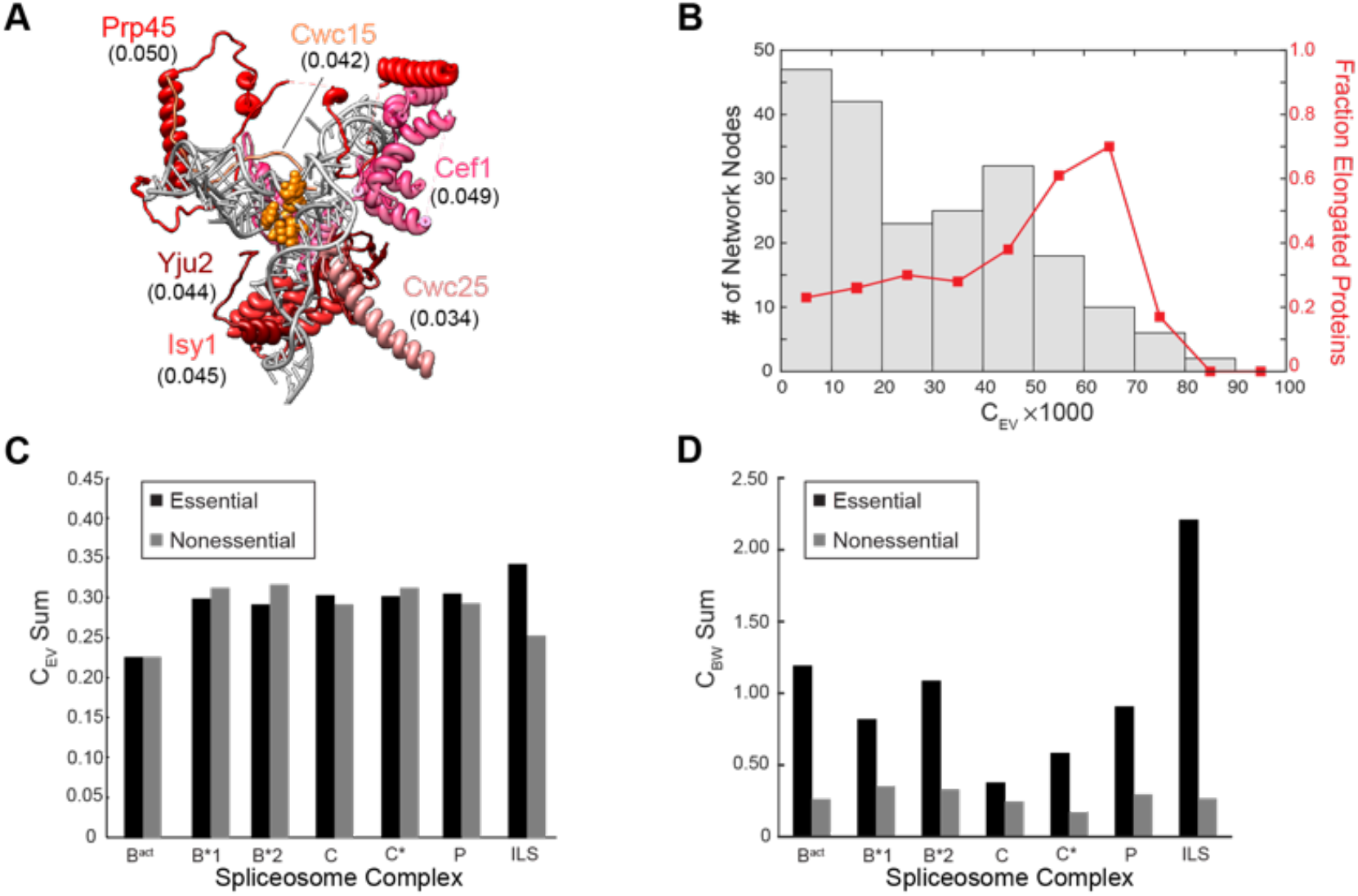
Contributions of Elongated and Nonessential Proteins to Spliceosome Network Connectivity. (**A**) View of the C complex spliceosome active site (PDB ID: 5LJ5) with elongated proteins with values of C_EV_ higher than the average for the complex (〈C_EV_〉= 0.032) are shown in shades of red and active site RNA nucleotides are shown in orange. C_EV_ values for the proteins are shown in parenthesis. (**B**) Histogram of the total number of network nodes in all spliceosome networks with given values of C_EV_ (grey) plotted in comparison with the fraction of those nodes originating from elongated proteins in each histogram bin (red). (**C**) Sum of C_EV_ values for essential and nonessential NTC/NTA components in each spliceosome structural network. (**D**) Sum of C_BW_ values for essential and nonessential NTC/NTA components in each spliceosome structural network.

We then examined how EF correlates with C_EV_ in spliceosome networks. We created a histogram in which we binned all of the calculated C_EV_ parameters for protein nodes. We then calculated the fraction of elongated proteins within each bin (**Fig. 5B**). The histogram reveals several features of the spliceosome networks. First, the distribution of C_EV_ is multi-modal. The largest bins correspond to splicing factors that are not highly interconnected and exhibit low C_EV_ values (0-0.03). A second cluster shows a peak within the 0.04-0.05 C_EV_ bin and tails off towards higher values of C_EV_. Bins containing proteins with C_EV_ >0.07 being dominated by Prp8. Thus, C_EV_ is not equally distributed across the nodes of spliceosome networks, rather these networks contain many nodes of both low and high influence.

Elongated proteins make up just over half of the nodes in the four highest C_EV_ bins (0.05<C_EV_<0.09; 53% elongated proteins; **Fig. 5B**). Furthermore, the vast majority of elongated proteins with high C_EV_ are members of the NTC/NTA complex. This indicates that elongated NTC/NTA proteins are the greatest protein influencers of spliceosome network architecture, surpassed as a group only by the globular protein Prp8.

Finally, we studied the relative contributions of essential and nonessential components of the NTC/NTA to the networks. Of 26 members of the NTC/NTA, eleven are nonessential for yeast growth under normal conditions while a twelfth (Cwc2) can be deleted if the yeast are grown at low temperatures (Hogg et al., 2014; Hogg et al., 2010). Despite being conditionally dispensable for splicing in yeast, nearly all of these nonessential proteins have human homologs. Surprisingly, the sum totals and averages for C_EV_ for nonessential components of the NTC/NTA was comparable, and in some cases, higher to those of the essential proteins (**Fig 5C**, **SI Appendix Table S7**). This shows that essentiality is not required for a factor to influence spliceosome network topology and that non-essential NTC/NTA components are collectively major contributors to network connectivity.

While non-essential components of the NTC/NTA are highly connected, they have low betweenness (C_BW_,**Fig. 5D**, **SI Appendix Table S8**) meaning that they do not form many unique bridges between other splicing factors. In contrast, essential components of the NTC/NTA have much higher C_BW_. Thus, while essential and non-essential components both have similar numbers of connections to highly connected splicing factors (C_EV_), only the essential components are likely to make unique connections between factors. Non-essential factors of the NTC/NTA may be able to modulate splicing activity due to the number of connections they make to central factors and yet may be dispensable due redundancy in these interactions.

## Discussion

Our network analysis has revealed general features of the splicing machinery derived from analysis of structural models representing different stages of the reaction. While spliceosomes are assembled from distinct subunits, the modularity of spliceosomes decreases significantly as the subunits integrate and the active site form (**Fig. 2A**). snRNAs remain in distinct modules following activation; however, other members of snRNA-containing modules frequently change (**Fig. 2B,C**). In concert with module dynamics, the connectivity of the spliceosome oscillates significantly during the reaction and is most connected during catalytic steps and least connected while transitioning to or from these structures (**Fig. 1B**). Thus, low connectivity networks link those of higher connectivity. While spliceosome composition changes dramatically during the reaction, changes in network topology are not explained by just the addition or subtraction of factors. Rather, rearrangements in the interactions between splicing factors are major contributors to the centrality parameters. Despite these dynamics, connectivity of the spliceosome is highly focused at the active site (**Fig. 4**). This is due in part to the role of Prp8 in scaffolding the large regions of the spliceosome together but also to the contributions of a number of elongated, non-globular proteins with high connectivity (**Fig. 5B**).

A recent study calculated network betweenness of spliceosome C complex factors and supported a well-established function of Prp8 as key communication conduit within spliceosome (Brow, 2002; Saltalamacchia et al., 2020). Our analysis provides further insights. First, we identify the U2 snRNA and Cef1 as additional key network nodes (**Fig. 3A**) due to their interactions with otherwise poorly-connected peripheral factors. Second, the network parameters vary from one structure to the next even for highly central factors like Prp8 and U2 (**Fig. 3B, C**). This supports the notion that key communication conduits of the spliceosome form transiently and can influence the network, and splicing, at specific stages. This also contrasts with conclusions made from networks derived from STRING database entries that provided evidence for limited impact of perturbations due to spliceosome modularity (Guimarães et al., 2018). Network perturbations, such as removal of a node corresponding to the Ecm2 protein (**SI Appendix Fig. S1B**), have consequences dependent on the structure of the spliceosome from which the network was calculated.

### Structural Networks Connect Betweenness with Lethality

Analysis of networks based on PPI databases (*e.g*., STRING) support a correlation between C_EV_ of network components and the lethality of the corresponding gene deletion (*i.e*., proteins with a large number of connections to other proteins are more likely to be essential for viability; the “centrality-lethality rule”) (He and Zhang, 2006; Jeong et al., 2001). This correlation was also evident in spliceosome networks derived from the STRING database (Guimarães et al., 2018). However, these types of networks fail to account for the dynamic nature of many interactions and how structural transitions can lead to transiently high C_BW_ for otherwise poorly connected factors. In our analysis of NTC/NTA components, we find essential and nonessential factors have similar C_EV_ but differing C_BW_ (**Fig. 5**). Thus, our work supports linking betweenness, or the uniqueness of connections rather than their numbers, with lethality. These results complement those obtained from networks derived from non-structural data and provide novel insights into relationships between network topologies and organism phenotypes.

### Transient, Disassortative Interactions Influence the Catalytic Core

Once the spliceosome has assembled and formed its active site, networks tended to show high C_EV_ and low C_BW_ for factors located near the active site (**Figs. 3**, **4**). The low C_BW_ values for nodes near the active site also suggests a mechanism by which spliceosome networks can be reversibly expanded or contracted. Based on network theory, splicing factors transiently bound to the spliceosome periphery can strongly influence network topology by providing more paths to one region or bridging components that were not previously connected. This results in asymmetrical or disassortative node interactions between poorly connected components and those of higher connectivity (Baingana and Giannakis, 2016; Newman, 2002). Networks with disassortative features, in turn, are particularly vulnerable to perturbation since incorporation of a single, peripheral factor can significantly impact the rest of network.

While disassortative spliceosome networks have previously been proposed based on biochemical and genetic interaction data (Pires et al., 2015), our work shows that such network features can also have a structural basis. Interactions between peripheral, poorly connected splicing factors with highly connected spliceosome core proteins, at specific steps, results in transient drops in algebraic connectivity (**Fig 1B**). Thus, the spliceosome network orients towards step-specific, peripheral factors, likely amplifying their influence on network communication. Notably, all of the spliceosome’s ATPases bind peripherally and exhibit disassortative interactions, consistent with their functions in altering spliceosome structure and influencing network topology (Staley and Guthrie, 1998; Yan et al., 2019). Whether or not regulators responsible for activating or repressing the splicing reaction would similarly impact spliceosome networks is not yet known.

### NTC/NTA Proteins are Highly Integrated and May Act as Regulatory Hubs

By comparison with splicing factors stably associated with the U snRNPs, several remarkable properties of the NTC/NTA proteins emerged from this network analysis. In the networks calculated here, the NTC/NTA never emerged as a distinct module. Instead, these factors dispersed throughout and transitioned between other modules (**Fig. 2**). This is consistent with a previous hypothesis based on functional assays in which the NTC/NTA no longer functions as a discrete complex after it has been integrated into the spliceosome (de Almeida and O’Keefe, 2015). The dispersion of the NTC/NTA is likely facilitated by the large number of NTC/NTA proteins which are elongated, rather than globular. Elongated peptide chains may permit formation of intermolecular contacts not possible with globular domains and this in turn leads to high C_EV_ for these components (**Fig. 5**). While it is possible that the NTC/NTA at some point can be segregated as a distinct module of spliceosome networks, it must involve a conformation of the spliceosome not captured by the models studied here.

Several components of the NTC/NTA are believed to help structure the spliceosome active site and modulate its activity (Hogg et al., 2014; Hogg et al., 2010; Rasche et al., 2012; Saha et al., 2012; van der Feltz et al., 2021; Villa and Guthrie, 2005). In the case of Cwc2, it has also been proposed that due to high degree of connectivity among spliceosomal proteins, Cwc2 is capable of transmitting local changes in structure to the RNA catalytic core to modulate activity (Rasche et al., 2012). This possible function may not be limited to Cwc2 since our data shows that many NTC/NTA proteins possess high degrees of connectivity (**Fig. 5**) and function as network nodes that bridge distal regions of the spliceosome as well as link the spliceosome’s core to its exterior. Unexpectedly, this is not limited to just essential NTC/NTA components as many nonessential factors have similar or even higher C_EV_. Since many NTC/NTA proteins are also elongated, their higher surface areas may facilitate increased numbers of interactions solely due to the increase in available binding sites. It is possible that this facilitates regulation of spliceosomal activity since it may maximize the impact of mutation, isoform expression, or post-translational modification of a single, highly-connected factor. The use of disordered regions to increase surface area available for protein-protein interactions may be an evolutionary solution shared between spliceosomal and ribosomal factors since almost half of ribosomal proteins also contain elongated domains (Peng et al., 2014).

Finally, network modeling and topological data such as C_EV_ and C_BW_ may be useful for inferring which NTC/NTA components are regulating splicing at distinct stages and the pathways by which that regulation may be occurring. For example, we identified the NTC/NTA factor Prp45 as having transiently high values of C_BW_ in B^act^ and ILS structures suggesting that it may be functioning as a regulatory node (**Fig. 3**). A regulatory function could result from Prp45 bridging factors that are unconnected in other structures. Prp45 has previously been shown to function in co-transcriptional spliceosome assembly, act in concert with the chromodomain protein Eaf3 to link chromatin state to spliceosome activation, and influence the recruitment of the Prp22 ATPase during the late stages of splicing (Gahura et al., 2009; Leung et al., 2019). Based on our networks, we would propose that Prp45 is regulating splicing during activation through its unique interactions with the RES complex and at later stages through similarly unique interactions with the NTR complex (**SI Appendix Fig. S2**).

## Conclusion

Our structure-based network analysis of spliceosomes highlights not only the utility of network modeling of an individual macromolecular structure but also the insights that can be obtained by comparison of networks representing successive stages along a reaction trajectory. As additional spliceosome structures emerge, it is likely that these methods can be extended for comparative analysis of spliceosomes present in different organisms (Fica, 2020). By reducing complex structures to network descriptors, this type of analysis may facilitate elucidation of the underlying interaction networks common to all spliceosomes and elaborated on by evolution.

## Materials and Methods

### Preparation of PDB Files for Network Analysis

Modeled structural coordinates for eight splicing complexes were downloaded from the RCSB PDB repository, PDB IDs: 5NRL, 5GM6, 6J6H, 6J6Q, 5LJ5, 5MQ0, 6EXN, and 5Y88. Multimeric Sm and Lsm rings as well as the Prp19 tetramer were treated as a single nodes in the network (**SI Appendix Table S1**). The pre-mRNA was split based on modeled regions into individual chains and nodes corresponding to the 5’ splice site, 5’ exon, branch site, 3’ exon, intron lariat, or mRNA as appropriate for each structure.

Chains identified as unknown were deleted or re-assigned based on their location, the electron density map, and/or comparison with independently determined structures (**SI Appendix Table S1**). In addition, a WD40 domain from a Prp19 monomer in C complex (PDB ID: 5LJ5) was reassigned to Prp17 based on the identical location of the protein domain in B*2 complex, where it is connected to Prp17.

In the B^act^ complex (PDB ID: 5GM6), nodes and edges were added to the network input parameters to connect the U2 snRNA with the U2 Sm ring, Lea1, and Msl1. In C* complex, edges were included in the network input parameters between Cef1 and Snt309 nodes as well as between the Cef1 and Prp19 nodes. In ILS complex (PDB ID: 5Y88), a portion of RNA could not be definitively modeled as either originating from the U6 snRNA or the intron and was therefore considered as its own unique node. Alanine-modeled regions were edited to remove interspersed glycines to permit analysis by PISA as described below. The total number of glycines removed from each structure is listed in **SI Appendix Table S1**.

### Spliceosome structural networks

Yeast spliceosomes structures were modeled as undirected networks, where RNA and proteins represent nodes, and intermolecular interactions among splicing factors define edges (Bastian et al., 2009). Using the edited PDB files, network edges were determined using PISA to identify any components with van der Waals surfaces closer together than 1.4 Å (Krissinel and Henrick, 2007). To automate these processes, we built custom software for extracting network information from PISA (LouiseNET, https://github.com/hoskinslab/LouiseNet, see **Supporting Materials and Methods**). LouiseNET can be used with any user-specified PDB file.

### Network Topology

Topological analysis of the spliceosome networks was performed using Gephi 0.9.2 (https://gephi.org). The global architectural features of the network, such as average degree, average clustering coefficient, and average path length were calculated using Gephi (Bastian et al., 2009). Algebraic connectivity is the second smallest eigenvalue of the Laplacian matrix and was calculated using custom Matlab scripts. Networks were visualized using the Yifan Hu proportional layout algorithm with default parameters in Gephi (Bastian et al., 2009).

### Modularity Analysis

Modularity analyses were performed on the adjacency matrix of each structure network, utilizing MODULAR (Marquitti et al., 2014). As previously described, a combination of a fast greedy and simulated annealing algorithms in MODULAR were used to calculate the modularity (Pires et al., 2015). The module distribution output from the fast greedy run was used as input for the simulated annealing algorithm. Protein and RNA assignments to different modules per structure were identified with MODULAR. Venn diagrams were generated with VennDiagram (Chen and Boutros, 2011).

### Centrality and Betweenness Analysis

Betweenness and eigenvector centralities were calculated using MatLab (MathWorks) following the formalized definitions found in (Newman, 2010). Eigenvector centrality was calculated following the formalized definitions found in (Newman, 2010). In order to compare centralities between networks, values for each network were normalized such that the sum of all centrality scores was 1 (Carley and Kim, 2008). Centrality values were mapped onto structures and displayed with UCSF Chimera (Pettersen et al., 2004).

## Acknowledgements

Thank you to Charles Schneider for help generating the Venn diagram in Fig. 2C. We thank Tim Grant and members of the Butcher and Brow laboratories for helpful discussions. We also thank Tim Grant, Charles Query, Samuel Butcher, Sierra Love, Pablo Alcón, and Terence Tang for comments on the manuscript.

