## Supplemental Information for "Network Theory Reveals Principles of Spliceosome Structure and Dynamics"

**This PDF file includes:**

Supporting Materials and Methods
Supporting Discussion S1
Supporting Results
Figures S1 to S5
Tables S1 to S8
Supporting References

### Supporting Materials and Methods

#### LouiseNet Software

LouiseNet is a Python-based application programming interface for the structural network analysis of the macromolecular assemblies such as spliceosome (https://github.com/hoskinslab/LouiseNet). LousieNet automates the process of extracting interactions among different components of a PDB file based upon their shared buried interfaces calculated by the PISA web portal (1). Next, LouiseNet performs the fundamental network analysis on the interactions generated from PISA. Briefly, an open-source web-based automation tool, Selenium (https://www.selenium.dev), and Google Chrome driver were used to automate the web navigations on PISA. First, the software uploads the user-specified PDB file on the PISA web portal. In the case of multiple PDB entries, files are uploaded one-by-one. The software then automatically uses the PISA web portal to calculate intermolecular interfaces. Once the output is displayed on the PISA webpage, beautifulsoup4 (https://pypi.org/project/beautifulsoup4/), a Python library for scraping data from HTML files, is used to extract the interface data. While not implemented in this work, LouiseNET can also use this same process to identify the interactions between individual residues of a chain with the rest of the chains in a PDB file. The software then saves the extracted output of the PISA webpage as a CSV file.

Next, we use a Pandas software library (https://pandas.pydata.org) to process the scraped data from PISA into a node and edge list format, which is subsequently used for network analysis. The edge list and node information are used to graph the network, calculate the topological network parameters (average degree, average clustering coefficient, and average shortest path length), and degree, eigenvector, and betweenness centralities for each node in the network. All the network calculations and graphs are calculated using the network analysis package, NetworkX (2), in Python. Additionally, LouiseNet provides users four different layouts to visualize the network. Finally, the network structure plot, network parameters, and centrality values are saved in a user-specified output folder.

**Supporting Discussion S1**

**Application of Network Theory to Cryo-EM Structures**

Our assumption in this analysis is that the model structures determined from the available cryo-EM data represent regions involved in stable interactions (or, alternatively, interactions observed in stable, non-catalytic structures) and key structural connections for the complex. It is likely that regions that are unresolved by the cryo-EM data represent flexible or structural dynamic domains. While these could play a role in spliceosome transitions, we cannot assign parameters or interactions to them.

Furthermore, basing the network nodes and interactions on whole RNA or protein chains both increases the computational feasibility of this study and avoids some biases that could arise from structural models built from cryo-EM data. Specifically, resolution is not consistent throughout spliceosome cryo-EM structures, making side chains on the periphery less likely to be resolved and correctly modeled in a specific conformation or interaction (3, 4). The robustness of our approach therefore stems from building our networks based on more reliable surface interactions between splicing factors that can be visualized over entire spliceosome complexes (e.g., Prp8 interacting with Snu114) rather than on atomistic details of those interactions (e.g., identification of specific amino acid or nucleotide contacts) that are only resolved in certain regions of the structure. Finally, we do not weight our network models based on surface areas of the modeled interactions since these values may have some error in low resolution regions of the structures and the significance of the surface area of an interaction may not necessarily be correlated with its impact on the splicing reaction.

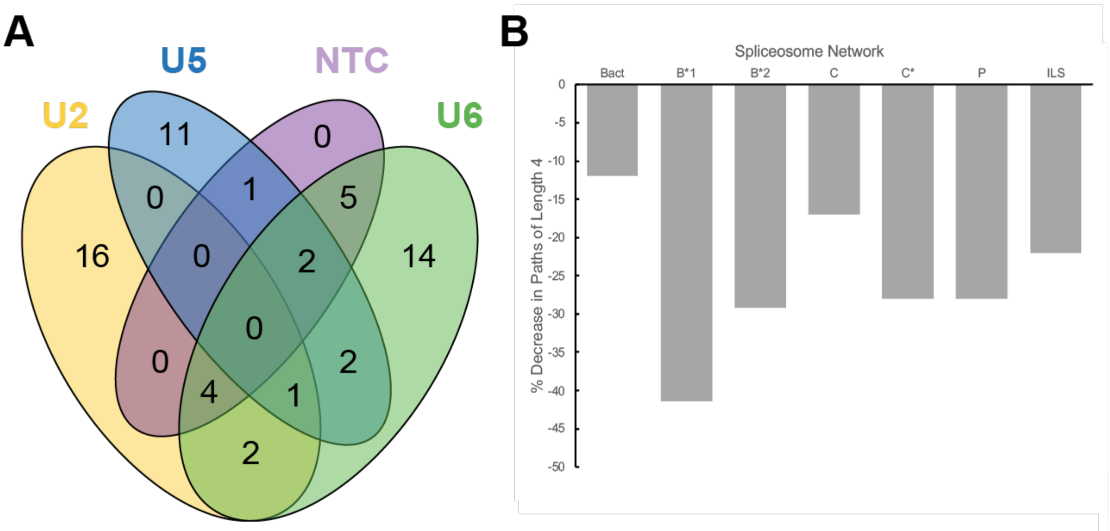

**Fig. S1. Consequences of Spliceosome Dynamics on NTC Module Association and Network Perturbations. (A)** Venn diagram of the module assignments for NTC/NTA factors over all spliceosome networks (B→ILS). (**B**) Impact of deletion of the node corresponding to the splicing factor Ecm2 on network algebraic connectivity (shown are the percent changes in the numbers of paths of length 4) for each spliceosome network.

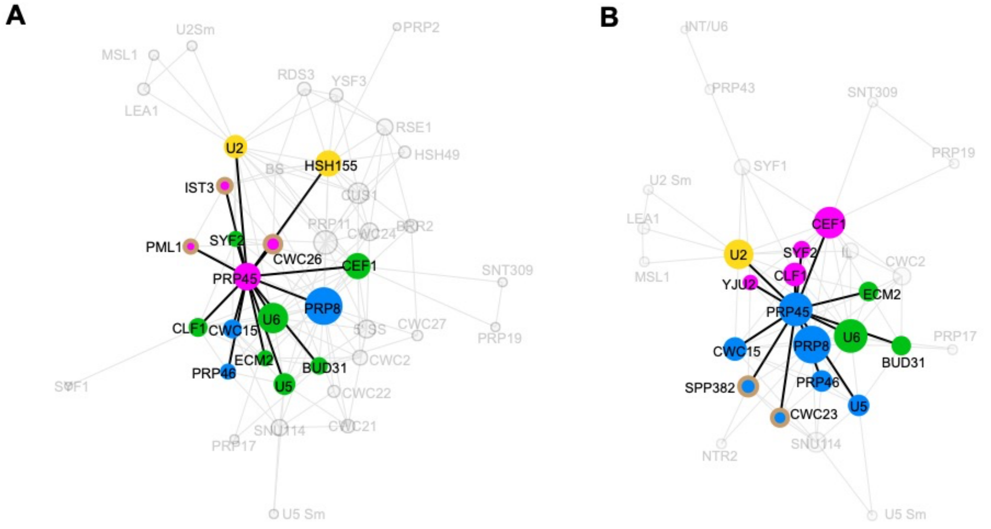

**Fig. S2. Prp45 connections in B$^{act}$ and ILS networks.** (**A**) Prp45 interaction network with other splicing factors in B$^{act}$ complex. Splicing factors are colored by module as shown in main text **Figure 2**. Interacting components of the RES complex are highlighted by circles with a tan border. (**B**) Prp45 interaction network with its partnering proteins in the ILS complex. Interacting components of the NTR complex are highlighted by circles with a tan border.

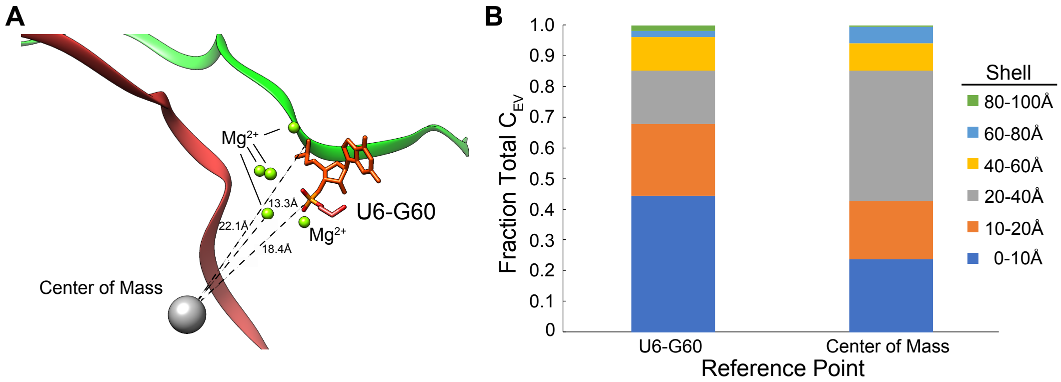

**Fig. S3. Eigenvector centrality is not always focused on the center of mass.** (**A**) Center of mass for the spliceosome B*2 model (PDB ID: 6J6Q) depicted in relation to the spliceosome active site (approximated as U6 nucleotide G60). The green ribbon represents the intron branch site and the red ribbon the 5' SS. (**B**) Sum of splicing factor $C_{EV}$ values as a function of node distance from the spliceosome active site U6-G60 or from the center of mass. Note that total $C_{EV}$ decreases in each shell when plotted as distance from the active site but not when plotted from the center of mass (compare the 20-40Å shells).

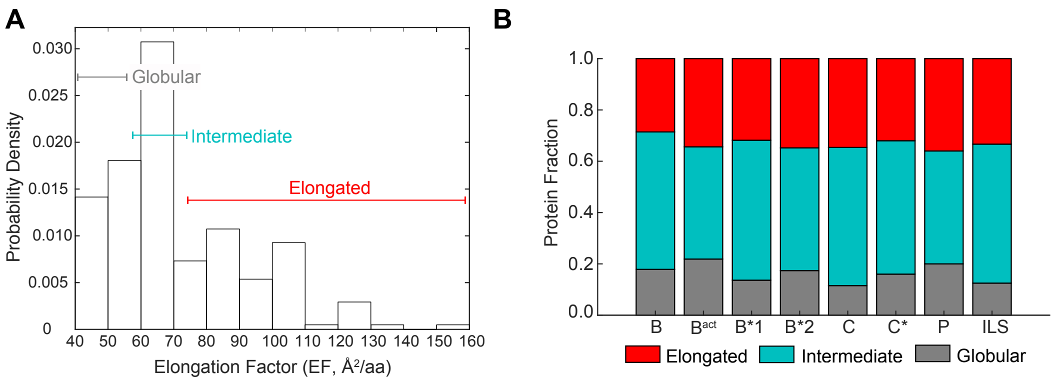

**Fig. S4. Identification of elongated, non-globular protein components of the spliceosome.** (**A**) The categorization per complex of splicing protein factors as elongated, intermediate, or globular based on chain surface area per amino acid (Å$^2$/aa). (**B**) Relative content of elongated, intermediate, and globular proteins found in each spliceosome structural model.

**Table S1.** Unresolved regions and glycine to alanine substitutions for each complex.

| | Complex | B | B^act | B*1 | B*2 | C | C* | P | ILS |
|---|---|---|---|---|---|---|---|---|---|
| **Structure** | PDB ID | 5NRL | 5GM6 | 6J6H | 6J6Q | 5LJ5 | 5MQ0 | 6EXN | 5Y88 |
| **Unresolved protein** | aa | 3855 | 6872 | 4542 | 4296 | 4891 | 5317 | 5633 | 5484 |
| | % | 22 | 33 | 34 | 32 | 29 | 35 | 38 | 38 |
| **Unresolved snRNA** | U2 nt | 1020 | 1109 | 966 | 965 | 1004 | 1020 | 1039 | 1093 |
| | % | 87 | 94 | 82 | 82 | 85 | 87 | 88 | 93 |
| | U4 nt | 46 | - | - | - | - | - | - | - |
| | % | 29 | - | - | - | - | - | - | - |
| | U5 nt | 44 | 97 | 35 | 35 | 73 | 73 | 43 | 97 |
| | % | 21 | 45 | 16 | 16 | 34 | 34 | 20 | 45 |
| | U6 nt | 17 | 5 | 4 | 4 | 14 | 8 | 9 | 6 |
| | % | 15 | 4 | 4 | 4 | 13 | 7 | 8 | 5 |
| **Substrate resolved** | nt | 29 | 75 | 60 | 57 | 50 | 53 | 71 | 28 |
| **Alanine modeling** | Chains affected | 0 | 0 | 2 | 2 | 8 | 10 | 6 | 1 |
| | Total gly-to-ala substitutions | 0 | 0 | 8 | 8 | 108 | 35 | 23 | 43 |
| | % substitutions | 0 | 0 | 3 | 3 | 2 | 1 | 2 | 6 |
| **Unknown chains** | Present | Yes | No | No | No | Yes | Yes | Yes | No |
| | Length (aa) | 30 | - | - | - | 132 | 68 | 95 | - |
| | Deleted | Yes | - | - | - | Partial | No | Yes | - |
| | Reassignment | - | - | - | - | 54 aa to Syf2 | all to Yju2 | - | - |
| **Grouped multimeric components** | LSm 2-8 | Yes | - | - | - | - | - | - | - |
| | U2 Sm B,D3,D1,D2,G,E,F | Yes | Yes | Yes | Yes | Yes | Yes | Yes | Yes |
| | U4 Sm B,D3,D1,D2,G,E,F | Yes | - | - | - | - | - | - | - |
| | U5 Sm B,D3,D1,D2,G,E,F | Yes | Yes | Yes | Yes | Yes | Yes | Yes | Yes |
| | Prp19 tetramer | - | Yes | Yes | Yes | Yes | Yes | Yes | Yes |

**Table S2.** Network topological parameters

| | Network | B | $B^{act}$ | B*1 | B*2 | C | C* | P | ILS |
|---|---|---|---|---|---|---|---|---|---|
| **Network descriptor** | Number of nodes | 34 | 40 | 27 | 28 | 31 | 31 | 30 | 29 |
| | Number of edges | 108 | 143 | 99 | 119 | 117 | 103 | 112 | 91 |
| | Average degree | 6.35 | 7.46 | 7.33 | 8.50 | 7.55 | 6.64 | 7.47 | 6.28 |
| | Avg. clustering coefficient | 0.68 | 0.63 | 0.70 | 0.71 | 0.71 | 0.65 | 0.64 | 0.68 |
| | Avg. path length | 2.36 | 2.18 | 1.98 | 1.83 | 1.92 | 2.08 | 1.96 | 2.21 |
| | Algebraic connectivity | 0.36 | 0.80 | 0.59 | 0.90 | 0.95 | 0.35 | 0.89 | 0.33 |

**Table S3A.** Components in U2 module per complex.

| B | $B^{act}$ | B*1 | B*2 | C | C* | P | ILS |
|---|---|---|---|---|---|---|---|
| U2 snRNA | U2 snRNA | U2 snRNA | U2 snRNA | U2 snRNA | U2 snRNA | U2 snRNA | U2 snRNA |
| U2Smring | U2Smring | U2Smring | U2Smring | U2Smring | U2Smring | U2Smring | U2Smring |
| LEA1 | LEA1 | LEA1 | LEA1 | LEA1 | LEA1 | LEA1 | LEA1 |
| MSL1 | MSL1 | MSL1 | MSL1 | MSL1 | MSL1 | MSL1 | MSL1 |
| HSH155 | HSH155 | SYF1 | SYF1 | | SYF1 | SYF1 | |
| RSE1 | RSE1 | ISY1 | | ISY1 | | | |
| BS | BS | BS | | LARIAT | | | |
| CUS1 | CUS1 | | | | | | |
| HSH49 | HSH49 | | | | | | |
| RDS3 | RDS3 | | | | | | |
| PRP9 | YSF3 | | | | | | |
| PRP11 | PRP11 | | | | | | |
| PRP21 | PRP2 | | | | | | |
| | CWC24 | | | | | | |
| | BRR2 | | | | | | |
| | | | | YJU2 | YJU2 | YJU2 | |
| | | | | CWC25 | | | |
| | | | | PRP17 | | | |
| | | | | | | SNT309 | |
| | | | | | | PRP19 | |
| | | | | | | CEF1 | |

*Note*: Grey Bar represent the absence of the corresponding protein in the respective complex.

**Table S3B.** Components in U6 module per complex.

| B | $B^{act}$ | B*1 | B*2 | C | C* | P | ILS |
|---|---|---|---|---|---|---|---|
| U6 snRNA | U6 snRNA | U6 snRNA | U6 snRNA | U6 snRNA | U6 snRNA | U6 snRNA | U6 snRNA |
| U4 snRNA | | | | | | | |
| Lsm2-8 ring | | | | | | | |
| PRP8 | | | | | | | |
| BRR2 | | | | | | | |
| DIB1 | | | | | | | |
| SNU66 | | | | | | | |
| PRP31 | | | | | | | |
| PRP3 | | | | | | | |
| PRP4 | | | | | | | |
| PRP6 | | | | | | | |
| SNU13 | | | | | | | |
| SNU23 | | | | | | | |
| PRP38 | | | | | | | |
| SPP381 | | | | | | | |
| U4Sm ring | | | | | | | |
| | CWC2 | CWC2 | CWC2 | | CWC2 | CWC2 | CWC2 |
| | BUD31 | BUD31 | BUD31 | | BUD31 | BUD31 | BUD31 |
| | PRP17 | PRP17 | PRP17 | | PRP17 | PRP17 | PRP17 |
| | | | | PRP45 | PRP45 | | |
| | ECM2 | | | | ECM2 | | ECM2 |
| | CEF1 | | | | CEF1 | | |
| | CLF1 | | | CLF1 | CLF1 | | |
| | SYF1 | | | SYF1 | | | |
| | SYF2 | | | SYF2 | SYF2 | | |
| | SNT309 | | | | SNT309 | | |
| | PRP19 | | | | PRP19 | | |
| | | | BS | | LARIAT | LARIAT | LARIAT |
| | | | YJU2 | | | | |
| | | | ISY1 | | | | |
| | | | 5'SS | | | | |
| | | | | PRP46 | | | |
| | | | | | | PRP18 | |

*Note*: Grey Bar represent the absence of the corresponding protein per complex.

**Table S3C.** Components in U5 module per complex.

| B | $B^{act}$ | B*1 | B*2 | C | C* | P | ILS |
|---|---|---|---|---|---|---|---|
| U5 snRNA | U5 snRNA | U5 snRNA | U5 snRNA | U5 snRNA | U5 snRNA | U5 snRNA | U5 snRNA |
| U5Sm ring | U5Sm ring | U5Sm ring | U5Sm ring | U5Sm ring | U5Sm ring | U5Sm ring | U5Sm ring |
| SNU114 | SNU114 | SNU114 | SNU114 | SNU114 | SNU114 | SNU114 | SNU114 |
| | PRP8 | PRP8 | PRP8 | PRP8 | PRP8 | PRP8 | PRP8 |
| | CWC22 | CWC22 | CWC22 | CWC22 | CWC22 | CWC22 | |
| | CWC21 | CWC21 | CWC21 | CWC21 | CWC21 | CWC21 | |
| | CWC15 | | | CWC15 | CWC15 | | CWC15 |
| | PRP46 | | | | PRP46 | | PRP46 |
| | CWC27 | | | | | | |
| | 5'SS | 5SS | | | | | |
| | | | | 5' EXON | 5' EXON | | |
| | | | | | | mRNA | |
| | | | | PRP16 | | | |
| | | | | BRR2 | | | |
| | | | | | SLU7 | SLU7 | |
| | | | | | PRP18 | | |
| | | | | | | PRP22 | |
| | | | | | | | PRP45 |
| | | | | | | | CWC23 |
| | | | | | | | SPP382 |
| | | | | | | | NTR2 |

*Note*: Grey Bar represent the absence of the corresponding protein per complex.

**Table S3D.** Components in non-snRNA module per complex.

| B | $B^{act}$ | B*1 | B*2 | C | C* | P | ILS |
|---|---|---|---|---|---|---|---|
| | PRP45 | PRP45 | PRP45 | | | PRP45 | |
| | PML1 | CEF1 | CEF1 | CEF1 | | | CEF1 |
| | IST3 | PRP19 | PRP19 | PRP19 | | | PRP19 |
| | CWC26 | SNT309 | SNT309 | SNT309 | | | SNT309 |
| | | CLF1 | CLF1 | | | CLF1 | CLF1 |
| | | SYF2 | SYF2 | | | SYF2 | SYF2 |
| | | ECM2 | ECM2 | ECM2 | | ECM2 | |
| | | CWC15 | CWC15 | | | CWC15 | |
| | | PRP46 | PRP46 | | | PRP46 | |
| | | | | CWC2 | | | |
| | | | | BUD31 | | | |
| | | | | | 3' EXON | | |
| | | | | | PRP22 | | |
| | | | | | | | SYF1 |
| | | | | | | | YJU2 |
| | | | | | | | PRP43 |
| | | | | | | | INT/U6 |

*Note*: Grey Bar represent the absence of the corresponding protein per complex.

**Table S4.** Degree and Betweenness differences between complexes for select components.

| Transition | B to $B^{act}$ | | $B^{act}$ to B*1 | | B*1 to B*2 | | B*2 to C | | C to C* | | C* to P | | P to ILS | |
|---|---|---|---|---|---|---|---|---|---|---|---|---|---|---|
| Component | ΔDeg | $\Delta C_{BW}$ | ΔDeg | $\Delta C_{BW}$ | ΔDeg | $\Delta C_{BW}$ | ΔDeg | $\Delta C_{BW}$ | ΔDeg | $\Delta C_{BW}$ | ΔDeg | $\Delta C_{BW}$ | ΔDeg | $\Delta C_{BW}$ |
| Prp8 | +9 | [green] | -5 | [grey] | +3 | [pink] | +3 | [green] | -2 | [green] | 0 | [pink] | -5 | [pink] |
| Prp45 | | | -4 | [pink] | +1 | [grey] | -1 | [grey] | 0 | [grey] | 0 | [grey] | +3 | [green] |
| Hsh155 | +6 | [green] | | | | | | | | | | | | |
| Cef1 | | | -1 | [green] | +2 | [pink] | -4 | [pink] | +1 | [green] | +3 | [green] | -2 | [grey] |
| Clf1 | | | +1 | [pink] | 0 | [grey] | -1 | [grey] | +2 | [green] | 0 | [grey] | -2 | [grey] |
| Cwc2 | | | +2 | [green] | +1 | [grey] | 0 | [grey] | -2 | [grey] | 0 | [grey] | -1 | [grey] |
| Syf1 | | | +6 | [green] | -1 | [grey] | -3 | [grey] | 0 | [grey] | +1 | [grey] | +1 | [green] |
| U6 snRNA | +3 | [pink] | -2 | [grey] | +3 | [pink] | 0 | [grey] | -3 | [pink] | +1 | [grey] | -2 | [green] |
| U2 snRNA | 0 | [pink] | +2 | [pink] | +4 | [grey] | 0 | [grey] | -6 | [grey] | +2 | [grey] | -2 | [green] |
| U5 snRNA | +5 | [green] | -1 | [grey] | +1 | [grey] | -1 | [grey] | 0 | [grey] | 0 | [grey] | -2 | [grey] |
| 5' Splice site | +6 | [grey] | +1 | [green] | +3 | [grey] | | | | | | | | |
| Branchsite | +2 | [grey] | -6 | [green] | +4 | [pink] | | | | | | | | |
| 5' exon | | | | | | | | | -2 | [grey] | | | | |
| Lariat | | | | | | | | | -3 | [grey] | +2 | [grey] | -4 | [grey] |

Note: Green/pink bars represent increase/decrease in $C_{BW}$ of a component above or below average value of the $C_{BW}$ of complex. Grey bar represent no change in $C_{BW}$ between consecutive splicing steps.

**Table S5.** Distances between the spliceosome active site and the structural center of mass

| | Complex | B | $B^{act}$ | B*1 | B*2 | C | C* | P | ILS |
|---|---|---|---|---|---|---|---|---|---|
| **Distance from active site (U6 G60) to center of structure in Å** | | 37.5 | 23.5 | 21.7 | 21.8 | 5.6 | 14.5 | 10.2 | 14.2 |

**Table S6.** Elongated proteins identified in each complex.

| B | B$^{act}$ | B*1 | B*2 | C | C* | P | ILS |
|---|---|---|---|---|---|---|---|
| SNU66 (91.7) | CEF1 (82.7) | CEF1 (82.8) | CEF1 (82.4) | CEF1 (81.4) | CEF1 (87.3) | CEF1 (80.2) | CEF1 (82.4) |
| PRP3 (80.7) | PRP45 (110.7) | PRP45 (106.2) | PRP45 (106.2) | PRP45 (108.8) | PRP45 (104.2) | PRP45 (109.6) | PRP45 (106.8) |
| SNU23 (91.8) | CWC15 (127.5) | CWC15 (128.2) | CWC15 (128.8) | CWC15 (133.8) | CWC15 (128.7) | CWC15 (125.9) | CWC15 (126.7) |
| | SYF2 (98.6) | SYF2 (99.6) | SYF2 (98.6) | ISY1 (83.0) | SYF2 (106.0) | SYF2 (105.2) | SYF2 (99.1) |
| | CUS1 (90.5) | ISY1 (155.6) | ISY1 (82.8) | YJU2 (81.8) | | | |
| | PRP11 (86.2) | | YJU2 (88.9) | CWC25 (87.5) | | | |
| | CWC24 (92.9) | | | | | | |
| SPP381 (76.2) | SNT309 (82.3) | SNT309 (81.7) | SNT309 (81.7) | SNT309 (75.6) | SNT309 (74.2) | YJU2 (78.1) | YJU2 (104.2) |
| CUS1 (85.9) | CWC21 (106.7) | CWC21 (106.9) | CWC21 (106.5) | CWC21 (76.1) | CWC21 (75.6) | SNT309 (75.6) | SNT309 (81.8) |
| PRP11 (76.4) | CWC26 (103.2) | | | | SLU7 (91.2) | CWC21 (79.6) | NTR2 (90.4) |
| RDS3 (84.2) | YSF3 (87.2) | | | | | SLU7 (103.5) | CWC23 (91.1) |
| Prp21 (100.2) | | | | | | PRP18 (103.6) | |

*Note*: Red colored proteins represent $C_{EV}$ above average and blue colored proteins represent $C_{EV}$ below average. Elongation factors are shown in parentheses.

**Table S7.** Average Cev for essential and nonessential NTC/NTA factors in each complex

| NTC/NTA Factors | $B^{act}$ | B*1 | B*2 | C | C* | P | ILS |
|---|---|---|---|---|---|---|---|
| Essential | 0.028 | 0.042 | 0.036 | 0.033 | 0.030 | 0.030 | 0.043 |
| Nonessential | 0.022 | 0.034 | 0.034 | 0.032 | 0.039 | 0.036 | 0.036 |

**Table S8.** Average $C_{BW}$ for essential and nonessential NTC/NTA factors in each complex

| NTC/NTA Factors | $B^{act}$ | B*1 | B*2 | C | C* | P | ILS |
|---|---|---|---|---|---|---|---|
| Essential | 0.149 | 0.117 | 0.136 | 0.042 | 0.058 | 0.091 | 0.276 |
| Nonessential | 0.026 | 0.039 | 0.037 | 0.027 | 0.021 | 0.037 | 0.038 |
